# Trends in *Escherichia coli* bloodstream infection, urinary tract infections and antibiotic susceptibilities in Oxfordshire, 1998-2016: an observational study

**DOI:** 10.1101/223107

**Authors:** Karina-Doris Vihta, Nicole Stoesser, Martin Llewelyn, T. Phuong Quan, Tim Davies, Nicola J. Fawcett, Laura Dunn, Katie Jeffrey, Chris Butler, Gail Hayward, Monique Andersson, Marcus Morgan, Sarah Oakley, Amy Mason, David H Wyllie, Derrick Crook, Mark H. Wilcox, Alan P. Johnson, Tim Peto, A. Sarah Walker

## Abstract

**Background:** The incidence of *Escherichia coli* bloodstream infections (EC-BSIs), particularly those caused by antibiotic-resistant strains, is increasing in the UK and internationally. This is a major public health concern but the evidence base to guide interventions is limited.

**Methods:** Incidence of EC-BSIs and *E. coli* urinary tract infections (EC-UTIs) in one UK region (Oxfordshire) were estimated from anonymised linked microbiological and hospital electronic health records, and modelled using negative binomial regression based on microbiological, clinical and healthcare exposure risk factors. Infection severity, 30-day allcause mortality, and community and hospital co-amoxiclav use were also investigated.

**Findings:** From 1998–2016, 5706 EC-BSIs occurred in 5215 patients, and 228376 EC-UTIs in 137075 patients. 1365(24%) EC-BSIs were nosocomial (onset >48h post-admission), 1863(33%) were community (>365 days post-discharge), 1346(24%) were quasi-community (31-365 days post-discharge), and 1132(20%) were quasi-nosocomial (≤30 days postdischarge). 1413(20%) EC-BSIs and 36270(13%) EC-UTIs were co-amoxiclav-resistant (41% and 30%, respectively, in 2016). Increases in EC-BSIs were driven by increases in community (10%/year (95% CI:7%–13%)) and quasi-community (8%/year (95% CI:7%–10%)) cases. Changes in EC-BSI-associated 30-day mortality were at most modest (p>0·03), and mortality was substantial (14-25% across groups). By contrast, co-amoxiclav-resistant EC-BSIs increased in all groups (by 11%-19%/year, significantly faster than susceptible EC-BSIs, p_heterogeneity_<0·001), as did co-amoxiclav-resistant EC-UTIs (by 13%-29%/year, p_heterogeneity_*0·001). Co-amoxiclav use in primary-care facilities was associated with subsequent co-amoxiclav-resistant EC-UTIs (p=0·03) and all EC-UTIs (p=0·002).

**Interpretation:** Current increases in EC-BSIs in Oxfordshire are primarily community-associated, with high rates of co-amoxiclav resistance, nevertheless not impacting mortality. Interventions should target primary-care facilities with high co-amoxiclav usage.

**Funding:** National Institute for Health Research.

**Research in context:** *Evidence before this study:* We searched PubMed for publications from inception up until October 26, 2017, with the terms *“Escherichia coli”, “E. coli”*, “bacteraemia”, “bloodstream infection”, restricting the search to English language articles, and also reviewed references from retrieved articles. *Escherichia coli (E. coli)* is the most common cause of bloodstream infection, and the incidence of *E. coli* bloodstream infection, and particularly antibiotic-resistant infections, is increasing in the UK and internationally. Although the UK government aims to reduce healthcare-associated *E. coli* bloodstream infection, there is only limited evidence to inform appropriate interventions.

**Added value of this study:** We investigated potential drivers for these increases in incidence by exploiting available linked electronic health records over 19 years for ~5200 patients with *E. coli* bloodstream infection and ~140000 with *E. coli* urinary tract infection, together with community antimicrobial prescribing data for the most recent six years. Our study identified several findings with significant implications for health policy and patient care:

- Increases in the incidence of *E. coli* bloodstream infections were driven mainly by non-hospital-associated cases; however, neither patients with previous urinary tract infections nor having previously had urine specimens sent from catheters appeared to be driving the increases
- Co-amoxiclav-resistant bloodstream infections rose significantly faster than co-amoxiclav-susceptible bloodstream infections, with the greatest number of co-amoxiclav-resistant bloodstream infections in 2016 being in patients discharged more than a month previously (i.e. community-associated)
- Higher co-amoxiclav use in primary care was associated with higher rates of both co-amoxiclav-resistant *E. coli* urinary tract infections and *E. coli* urinary tract infections overall, supporting drives to reduce broad-spectrum and inappropriate antibiotic use in primary care
- Despite substantial increases in co-amoxiclav-resistant bloodstream infections there was no evidence that mortality was increasing in these cases; this does not support moving to broader empiric antibiotic prescribing in hospitals (i.e. carbapenems, piperacillin-tazobactam)

**Implications of all available advice:** This suggests that government strategies to effectively reduce *E. coli* bloodstream infections should target community settings, as well as healthcare-associated settings. The absence of an increased mortality signal suggests that co-amoxiclav resistant *E. coli* infections are either being successfully treated by dual empiric therapy in severe cases (e.g. with concomitant gentamicin), can be “rescued” once isolate susceptibilities become available, or currently deployed phenotypic susceptibility testing breakpoints do not adequately correlate with clinical outcome.

## Introduction

*Escherichia coli* is a major cause of bloodstream infection (BSI)^1^ and a critical antimicrobial resistance (AMR) concern;^2^ rates are rising worldwide.^3^ For example, *E. coli* bloodstream infections (EC-BSIs) reported (voluntarily) to Public Health England rose by 44% between 2003-2011;^4^ a similar 68% increase between 1999-2011 was seen in Oxfordshire, UK.^5^ Mandatory reporting was introduced in England in July 2011; a further 28% increase in EC-BSI incidence occurred by July-September 2016, to 78·8 cases/100,000 population.^6^

In the UK, as elsewhere, most (>70%) EC-BSIs are identified within two days of hospital admission.^6^ However, the impact of previous hospital-exposure on trends in EC-BSI has not been comprehensively investigated, with only two relevant previous studies, one in the Calgary Health Region 2000-2006^7^, and another in Oxfordshire in 2011^5^ considering only whether blood cultures were taken outside or inside hospital. EC-BSI source may also differ by hospital-exposure. In a recent study, ~50% of UK EC-BSIs were considered most likely due to urinary tract infections (UTIs);^8^ gastrointestinal foci are however more common in inpatients.^6^

30-day all-cause mortality following EC-BSI is ~16%;^9^ and could rise given the impact of increasing AMR on treatment options.^2^ In Oxfordshire, EC-BSI incidence rises through 2011 were essentially confined to ciprofloxacin-, co-amoxiclav-, cefotaxime- and/or aminoglycoside-resistant organisms.^5^ The reasons for rising EC-BSI more generally are unclear, with increased antibiotic usage implicated in some, but not all, studies.^10–15^ In the UK and internationally, co-amoxiclav is used as empiric treatment for many infection syndromes and for prophylaxis.^10,16^ Hence, trends in co-amoxiclav resistance are particularly important.

We therefore aimed to investigate possible drivers of changes in EC-BSI incidence and antibiotic susceptibilities in Oxfordshire over the last two decades, while stratifying for hospital-exposure. We hypothesized that increases may be due to features of the at-risk population (therefore exploring demographics, recurrent infections, increased ascertainment), healthcare-history (previous urine cultures, and specifically previous catheter specimens, previous admission diagnoses, antibiotic usage), and/or the bacteria (exploring mortality/severity, AMR burden).

## Methods

The Infections in Oxfordshire Research Database (IORD)^17^ records all admissions to the Oxford University Hospitals National Health Service Foundation Trust (OUH), Oxfordshire, UK, from April 1997, linked by patient with microbiology and biochemistry/haematology results. The four hospitals within OUH provide all acute care, microbiology and pathology services in the region (~680,000 individuals). Out-of-hospital mortality was determined by updates from a national information system recording all UK deaths. IORD has generic Research Ethics Committee and Health Research Authority approvals (14/SC/1069, ECC5-017(A)/2009). Data on antibiotic prescribing and numbers of registered patients for each general practice were obtained from the Health and Social Care Information Centre (available January 2011-December 2016 only).

The primary study outcome was EC-BSI, defined as *E. coli* isolated from blood cultures taken 01/Jan/1998-31/Dec/2016 inclusive, including polymicrobial cultures (13%), without age restriction and de-duplicated within 14-days of each index positive.^18^ For context we also analysed *E. coli* UTIs (EC-UTIs), defined as pure culture from urine of >10^4^ colony-forming-units/ml, de-duplicated within 90-days. We classified EC-BSIs/EC-UTIs as ‘nosocomial’ if samples were taken >48h post-admission until discharge.^19^ All other EC-BSIs/EC-UTIs were classified as ‘community’, ‘quasi-community’ or ‘quasi-nosocomial’ if the last hospital discharge was >1 year, 31-365 days, or 0-30 days previously. We also calculated incidences of first ever and recurrent EC-BSIs. See Supplementary Methods for further details.

To account for the contribution of ageing and population growth, we standardised incidence for age and sex against the 1998 Oxfordshire population distribution (estimates from the UK Office for National Statistics). To assess ascertainment, we considered the incidence of blood/urine cultures, regardless of result, and also additionally standardised for culture rates. As a proxy for changes in bacterial virulence, we considered 30-day mortality after, and levels of monocytes, neutrophils, lymphocytes, C-reactive protein (CRP), creatinine and urea at (closest value within [−2,+2] days) sample collection. To investigate AMR burden, which might also affect treatment outcomes, we assessed resistance to drugs consistently tested throughout the study period. Susceptibility testing was performed using disk-diffusion to 31/Jan/2013, then by microbroth dilution (BD Phoenix™ Automated Microbiology System, Beckton Dickinson, Franklin Lakes, NJ, USA) (see Supplementary Methods).

Guidelines recommend empirical treatment for uncomplicated UTIs and for urine samples to be sent to the laboratory only from individuals with clinical treatment failure, frequent or recurrent UTI or with a possibly resistant infection.^16^ To investigate this patient group, we first classified EC-BSIs according to whether the patient had ever had an EC-UTI identified by the laboratory ≥3 days previously. To investigate the contribution of UTI around the time of the EC-BSI, including where *E. coli* was not isolated, we classified EC-BSIs as ‘likely urine-associated’ (urine sample taken 3-30 days previously; EC-UTI or mixed growth/negative but UTI suspected clinically from request codes), ‘urosepsis’ (defined as for likely urine-associated BSIs but urine samples within (−3,+2] days of the EC-BSI), ‘unlikely urine-associated’ (UTI with non-E. *coli* pathogen or no urine sample), or ‘unknown’ (other) (details in Supplementary Methods). To investigate the contribution of catheters, we classified EC-BSIs according to whether the patient had ever had a catheter urine specimen submitted up to and including the day of blood collection (regardless of result).

To investigate the contribution of previous admission characteristics, we classified quasi-nosocomial EC-BSIs by whether the primary diagnostic code of the antecedent admission was infection-related, or any diagnostic code (primary/secondary) included UTI (Supplementary Methods).

### Statistical analysis

Incidence was modelled using negative binomial regression of counts per month, binary outcomes using poisson regression of monthly counts (to estimate analogous rate ratios) and test results using median quantile regression of absolute values against sample date. Test results and mortality were adjusted for age and sex. Changes in trends in these outcomes were estimated using iterative sequential regression (Supplementary Methods),^20^ and compared between outcomes using stacked regression.^21^ To estimate associations with primary care co-amoxiclav prescribing, co-amoxiclav defined-daily-doses (DDDs) per 1000 registered patients in the previous or current year and general practice were included as explanatory variables (Supplementary Methods).

Analyses were conducted using R 3.2.2, and STATA 14.1 for stacked regression and probability weighted analyses.

## Results

After 14-day de-duplication, from 1998-2016 5706 EC-BSIs occurred in 5215 patients (i.e. 9% recurrences (relapse and/or reinfection)). Recurrences occurred a median(IQR) 144(39-577) days apart: of 391 patients with recurrences, 324(83%) had one and 52(13%) had two (range 1-8). Overall incidence increased year-on-year (annual incidence rate ratio (IRR)=1 06 (95% CI 1·05-1·06)). For most EC-BSI (5393(95%)) patients were admitted to OUH before or within the 24h following the blood culture (remainder mostly taken in emergency departments or community hospitals). Only 1365(24%) EC-BSIs were ‘nosocomial’ (≥48h post-admission). A further 1132(20%) were ‘quasi-nosocomial’ (discharged up to 30 days previously), 1346(24%) were ‘quasi-community’ (discharged 31365 days previously) and 1863(33%) were ‘community’ cases (discharged >1 year previously or never previously admitted to OUH).

Incidence trends for EC-BSIs varied substantially with hospital-exposure (**Figures 1A&2A, Supplementary Table 1**), with overall increases clearly driven by community and quasicommunity hospital-exposure groups, and no evidence of different incidence trends between these two groups in 2016 (p_heterogeneity_=0·27). By contrast, quasi-nosocomial and nosocomial EC-BSIs increased more slowly. Considering only the first EC-BSI per patient or subsequent EC-BSIs (**Figure 2A, Supplementary Figure 1**) gave broadly similar results. Year-on-year increases in first EC-BSI became smaller (but still significant) the more recent the hospital exposure. Quasi-community recurrent EC-BSI were rising faster than first EC-BSIs (p_heterogeneity_<0·001) and the stable current trend in all quasi-nosocomial BSIs appeared to be driven by reduced recurrences in this group.

**Figure 1.**
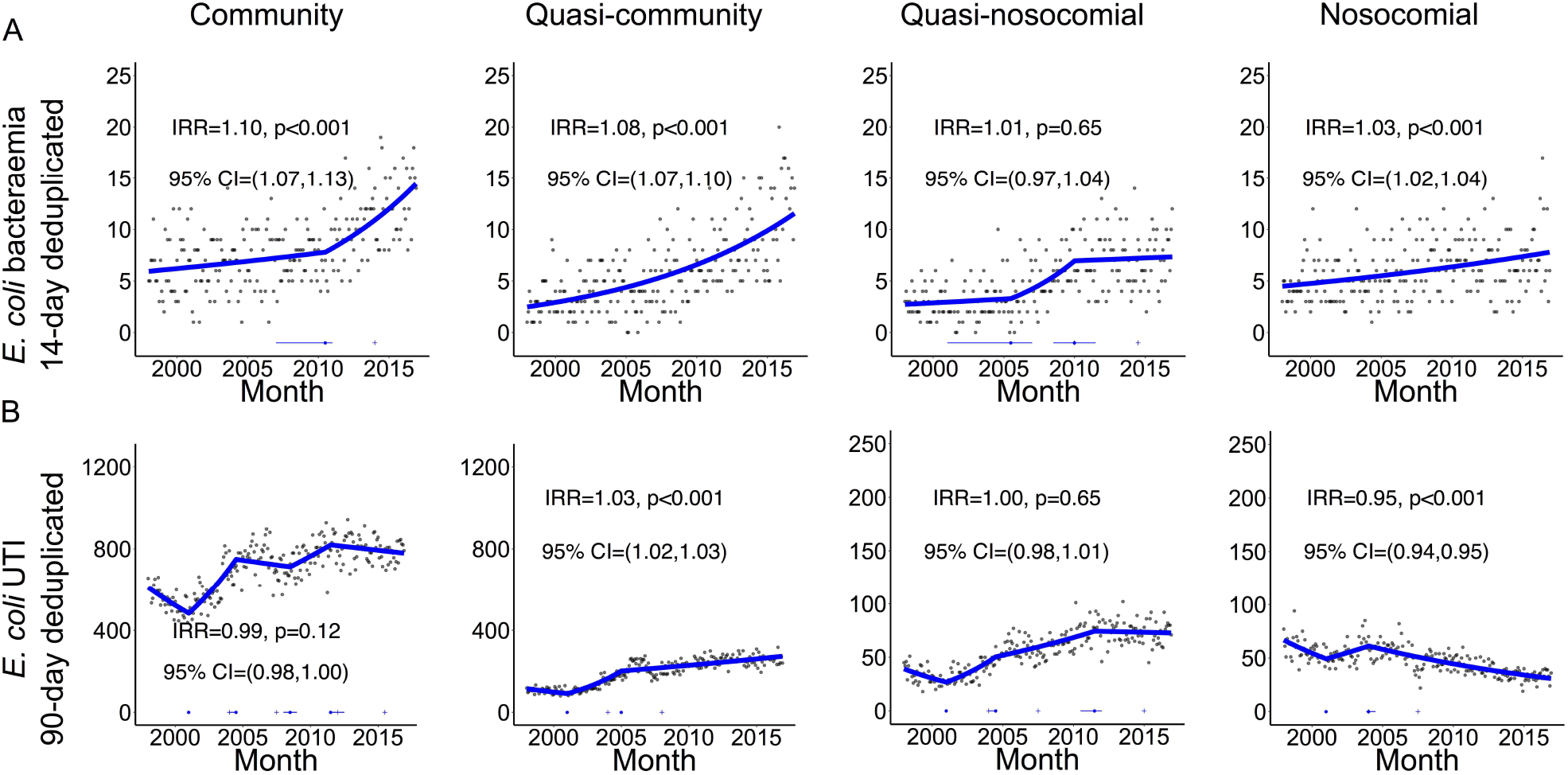
Monthly (A) EC-BSI and (B) EC-UTI according to recent hospital-exposure (first and recurrent infections). Footnote: only counting EC-BSI recurrences occurring >14 days after an index positive, and EC-UTI recurrences occurring >90 days after an index positive. Thick blue line represents the estimated incidence by iterative sequential regression (ISR). Blue lines at the base of the graph represent 95% CI around the breakpoints estimated by the ISR model. IRR=annual incidence rate ratio in 2016

**Figure 2.**
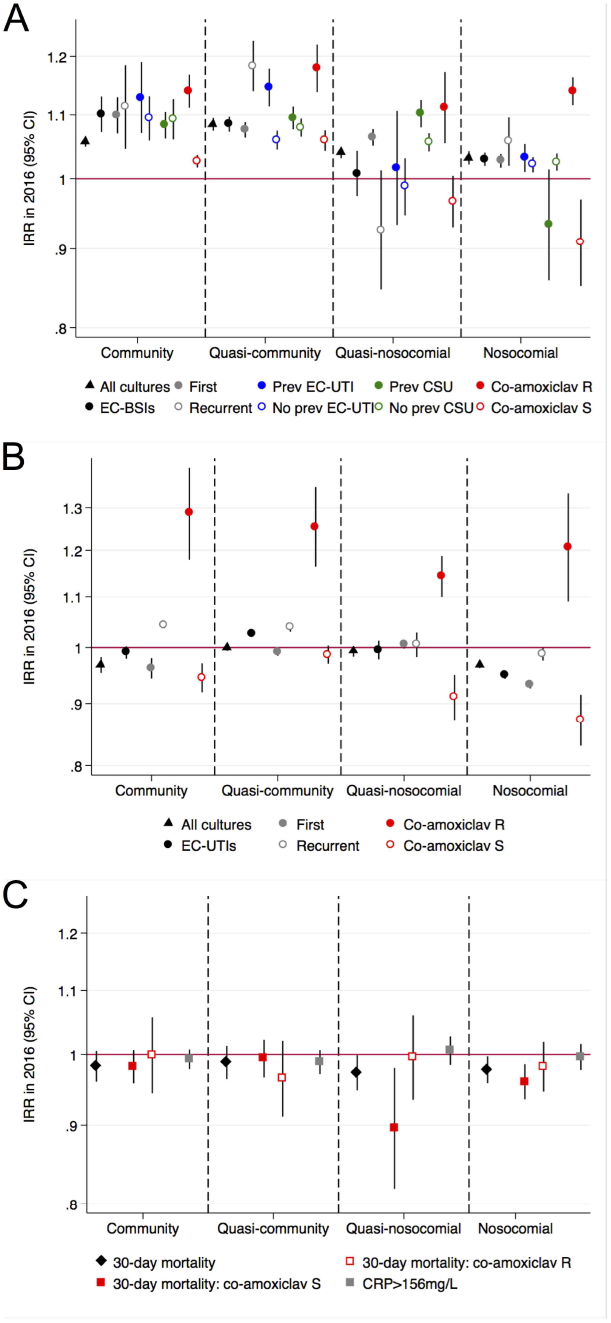
Summary of incidence trends in 2016 for (A) EC-BSIs, (B) EC-UTIs, and (C) severity of co-amoxiclav resistant and sensitive EC-BSIs. Footnote: IRR=annual incidence rate ratio in 2016. See Supplementary Table 1 for numbers and heterogeneity tests

After 90-day de-duplication, 228376 EC-UTIs occurred in 137075 patients (i.e. 40% recurrences (relapse/re-infection)). Recurrences occurred a median(IQR) 457(200-1119) days apart: of the 41371(30%) patients with recurrences, 22011(53%) had one and 8742(21%) had two (range 1-33). 12898(9%) patients had two EC-UTI within six months. EC-UTIs were predominantly community (160359,70%), and less commonly quasicommunity (44283,19%), quasi-nosocomial (12764,6%) or nosocomial (10970,5%) in origin. Rates of EC-UTI increased over 1998-2016 in community, quasi-community and quasi-nosocomial groups, although current trends were fairly stable, but declined significantly in the nosocomial group (**Figure 1B&2B**). Furthermore, increases were accounted for entirely by substantial increases in recurrent UTI episodes, with decreasing overall trends in first EC-UTI per patient (**Supplementary Figure 2**).

In 2016, therefore, recurrences accounted for at least half of community, quasi-community and quasi-nosocomial EC-UTIs, and around a fifth of quasi-community and quasi-nosocomial EC-BSIs (**Supplementary Table 2**).

### Impact of population and sampling on EC-BSI

Blood culture submission rates increased substantially from 1998-2016 for community/quasi-community/quasi-nosocomial groups (**Figure 2A, Supplementary Figure 3**), raising the possibility that observed increases in EC-BSIs were driven by increases in the use of blood cultures as a diagnostic test. However, there was no suggestion that the indications for blood culture changed with time: neutrophils and CRP when cultures were taken did not meaningfully change and there was no change in the 30-day mortality post blood culture sampling (**Supplementary Figure 4**). Further, increases in community blood culture submission rates were significantly smaller than increases in community EC-BSIs (p<0·001, **Figure 2A**). Standardising for age and sex explained only 10-26%, and standardising additionally for number of blood cultures taken 9-28%, of the increase in overall or first-per-patient EC-BSIs, with the greatest percentage explained in nosocomial EC-BSIs and the least in community EC-BSIs (**Supplementary Tables 3,4**). In contrast, urine sample submission was more stable over time (**Supplementary Figure 5**).

### Disease severity of EC-BSIs

30-day mortality following EC-BSI declined slightly (IRR=0·98) in the nosocomial (p=0·03) and quasi-nosocomial (p=0·06) groups, but there was no evidence for changes in quasicommunity and community groups (p>0·21, adjusting for age and sex, **Supplementary Figure 6**). Mortality was substantial at 25%, 30%, 16% and 14% across the groups, respectively. Changes in haematology/biochemistry test results over time were small and/or non-significant (**Supplementary Figure 6**), and did not indicate that less severe infections were being identified, or that there were any changes in pathogen virulence.

### Impact of previous illness on EC-BSI

1755(31%) EC-BSI occurred in patients with an EC-UTI ≥3 days previously (median(IQR) 213(43-918) days previously). However, incidence trends were broadly similar for EC-BSIs with or without EC-UTIs ≥3 days previously, although quasi-community EC-BSIs were rising particularly fast in those with previous EC-UTIs (p_heterogeneity_<0·001, **Figure 2A, Supplementary Figure 7**). We next explored whether EC-BSI increases were associated with past symptomatic urinary disease, including those without positive urine cultures. Considering urine samples/results taken within 30 days before the EC-BSI, and incorporating information on mixed growth and request codes, only 760(13%) EC-BSIs were ‘likely urine-associated’, with 1613(28%) ‘urosepsis’, 1613(28%) ‘unlikely urine-associated’ (of which 181[11%] had a contemporaneous urine specimen positive for another pathogen), and 1720(30%) unknown. However, the relative proportions of these did not vary substantially over time (**Figure 3**), suggesting no specific subgroup was associated with incidence increases. Percentages of EC-BSIs with a previous catheter urine specimen (CSU) increased with recency of hospital-exposure, being present in 365(20%) community, 364(32%) quasi-community, 541(40%), quasi-nosocomial, 584(43%) nosocomial. However, incidence trends were broadly similar for EC-BSIs with or without a previous CSU (**Figure 2A, Supplementary Figure 8**), although quasi-nosocomial EC-BSIs were rising particularly fast in those with previous CSUs (p_heterogeneity_<0·001), while increases in nosocomial EC-BSIs were restricted to those without previous CSUs (p_heterogeneity_=0·03).

**Figure 3.**
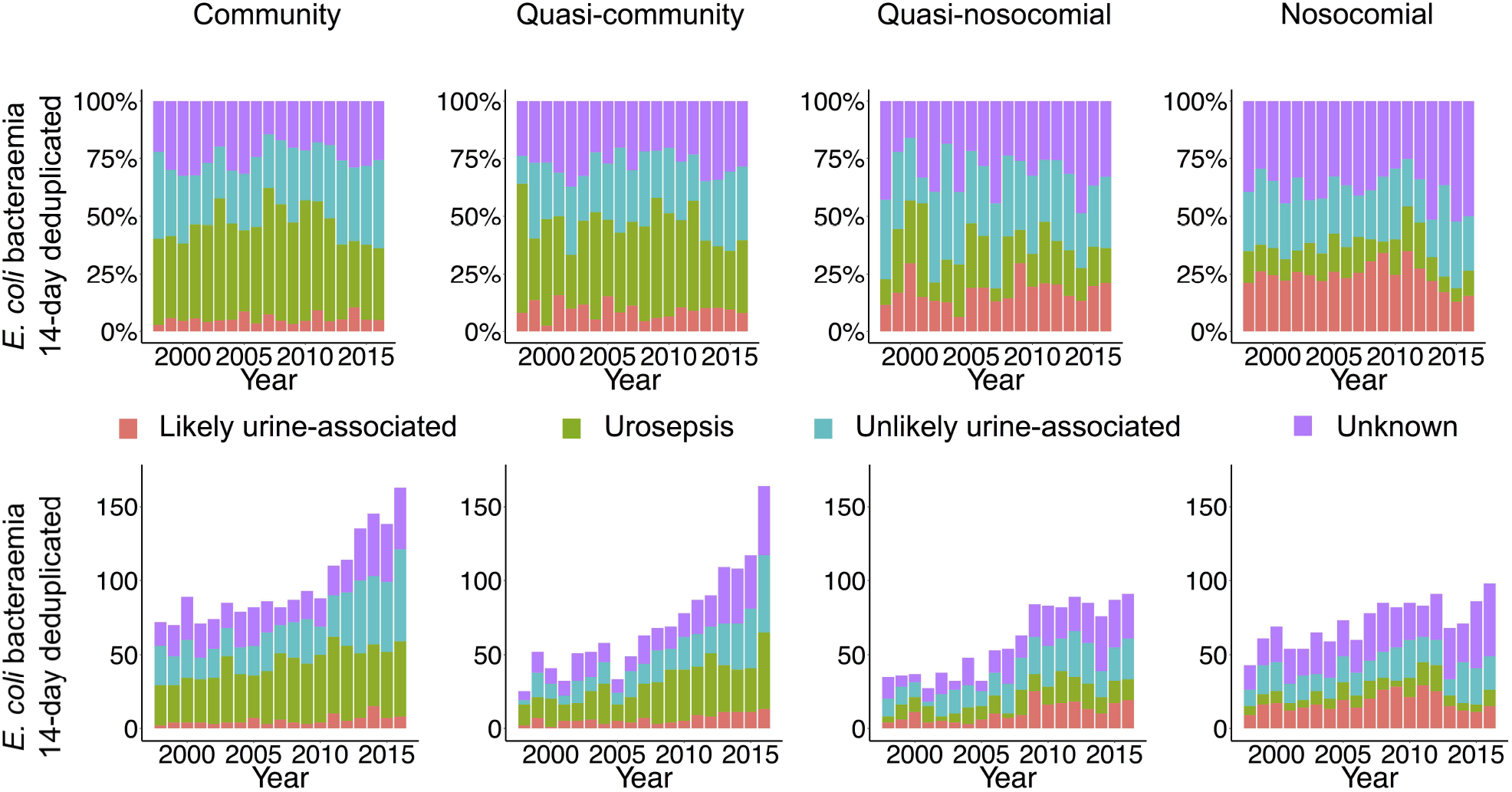
Annual EC-BSI according to recent hospital-exposure and urine sample submission/results. Footnote: See Supplementary Methods for definitions.

For the 1132 quasi-nosocomial EC-BSI patients discharged in the preceding 30 days, the most common reasons for the antecedent admission were malignancy (395,35%), gastrointestinal disorders (177,16%), and renal/urological disorders (164,14%) (**Supplementary Table 5**), with no major temporal variability (**Supplementary Figure 9A**). There was no evidence that the antecedent admission was shorter than the quasicommunity group (median 2·0 (IQR:0·3-7·9) days vs 23 (0·3-8·2) respectively, ranksum p=0·15). There was strong evidence that quasi-nosocomial EC-BSIs with a UTI diagnostic code or an infectious primary diagnostic code for the antecendent admission were rising faster than those without (heterogeneity p=0·005, p<0·001 respectively, **Supplementary Figure 9B&C**), but these still comprised <25% of quasi-nosocomial EC-BSIs.

### Antimicrobial susceptibility

Exploring the possibility that EC-BSI increases were associated with the development of AMR, the only EC-BSI antibiotic-resistant phenotype that consistently increased across all groups was co-amoxiclav (p<0·001; **Figures 2A&4**), with 212(41%) of 515 EC-BSIs in 2016 being co-amoxclav resistant (**Supplementary Table 6**). Co-amoxiclav-resistant EC-BSIs increased significantly faster than co-amoxiclav-susceptible EC-BSIs (p_heterogeneity_<0·001), but community and quasi-community co-amoxiclav-susceptible EC-BSIs were still increasing significantly in 2016. Most (942/1412, 67%) co-amoxiclav-resistant EC-BSIs remained susceptible to gentamicin and ciprofloxacin (**Figure 4**).

**Figure 4.**
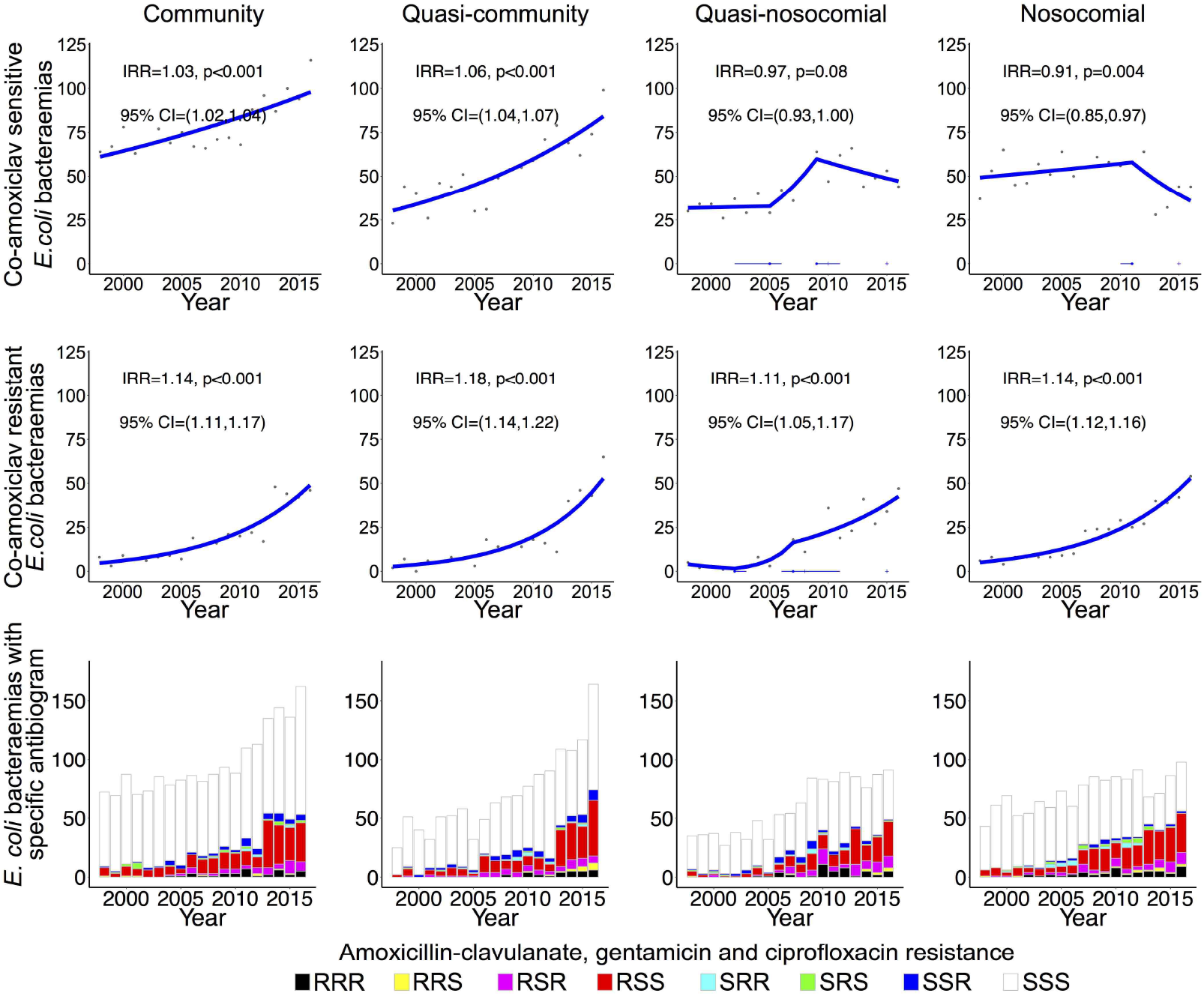
Annual EC-BSI susceptible and resistant to co-amoxiclav, with and without resistance to gentamicin and ciprofloxacin, according to recent hospital-exposure. Footnote: IRR=annual incidence rate ratio in 2016.

Increases in other antibiotic-resistant EC-BSIs were most notable in the community and quasi-community groups, with significant year-on-year increments in all but trimethoprim-resistant EC-BSIs, which remained stable in these groups (**Supplementary Figure 10**). Co-amoxiclav-resistant EC-UTIs also rose consistently and significantly regardless of healthcare-exposure, but trends were more variable for other antibiotics (**Supplementary Figure 11**). In 2016, 3921/13792(28%) EC-UTIs were co-amoxiclav-resistant.

Given the substantial increase in co-amoxiclav resistant EC-BSIs, we investigated whether there was any evidence of differential severity in susceptible and resistant cases. There was no strong evidence that co-amoxiclav-resistant EC-BSIs were associated with higher neutrophil counts in any hospital-exposure group (p>0·04, adjusting for age and sex), or that neutrophil counts were changing differently over time compared with co-amoxiclav-susceptible EC-BSI (p_heterogeneity_>0·67; **Supplementary Figure 12**). Mortality was higher (32% (95% CI 13%–46%); p=0·002, adjusting for age and sex) for co-amoxiclav-resistant vs co-amoxiclav-susceptible nosocomial EC-BSIs, but not community/quasi-community/quasi-nosocomial EC-BSIs (p>0·48), and mortality did not change differently over time in any group (p_heterogeneity_>0·35; **Supplementary Figure 12, Figure 2C**).

Over financial years 2003-2014, the strongest associations with nosocomial co-amoxiclav-resistant EC-BSIs were with hospital co-amoxiclav (cross-correlation 075) and third-generation cephalosporin (0·80) use (**Supplementary Table 7**). Community prescribing data was only available from 2011, and co-amoxiclav-resistant EC-BSIs were too few to consider relationships with co-amoxiclav use. However, from 2012-2016, primary care facilities prescribing more co-amoxiclav in the previous year had higher rates of subsequent co-amoxiclav-resistant-community-EC-UTIs (IRR (per 100DDD higher)=105 (95% CI 102108) p=0·003, **Figure 5**), and co-amoxiclav use in the previous year was a stronger predictor of current rates than co-amoxiclav use in the current year (p=0.003 vs p=0.64). Co-amoxiclav use in the current year was a stronger predictor of all community-EC-UTIs (p=0.01 vs p=0.11) and urine specimen submission (p=0.0001 vs p=0.006), and was associated with higher rates of both (IRR=1·02 (1·00-1·04) p=0·01 and 1.02 (1·01-1·03) p=0 002 respectively). Co-amoxiclav use was not associated with the proportions of *E. coli*-positive specimens (p=0·68). Similar results were seen across all samples regardless of hospital-exposure group (**Supplementary Figure 13**), and also when adjusting instead for the proportion aged over 65 and male in 2017 per practice.

**Figure 5.**
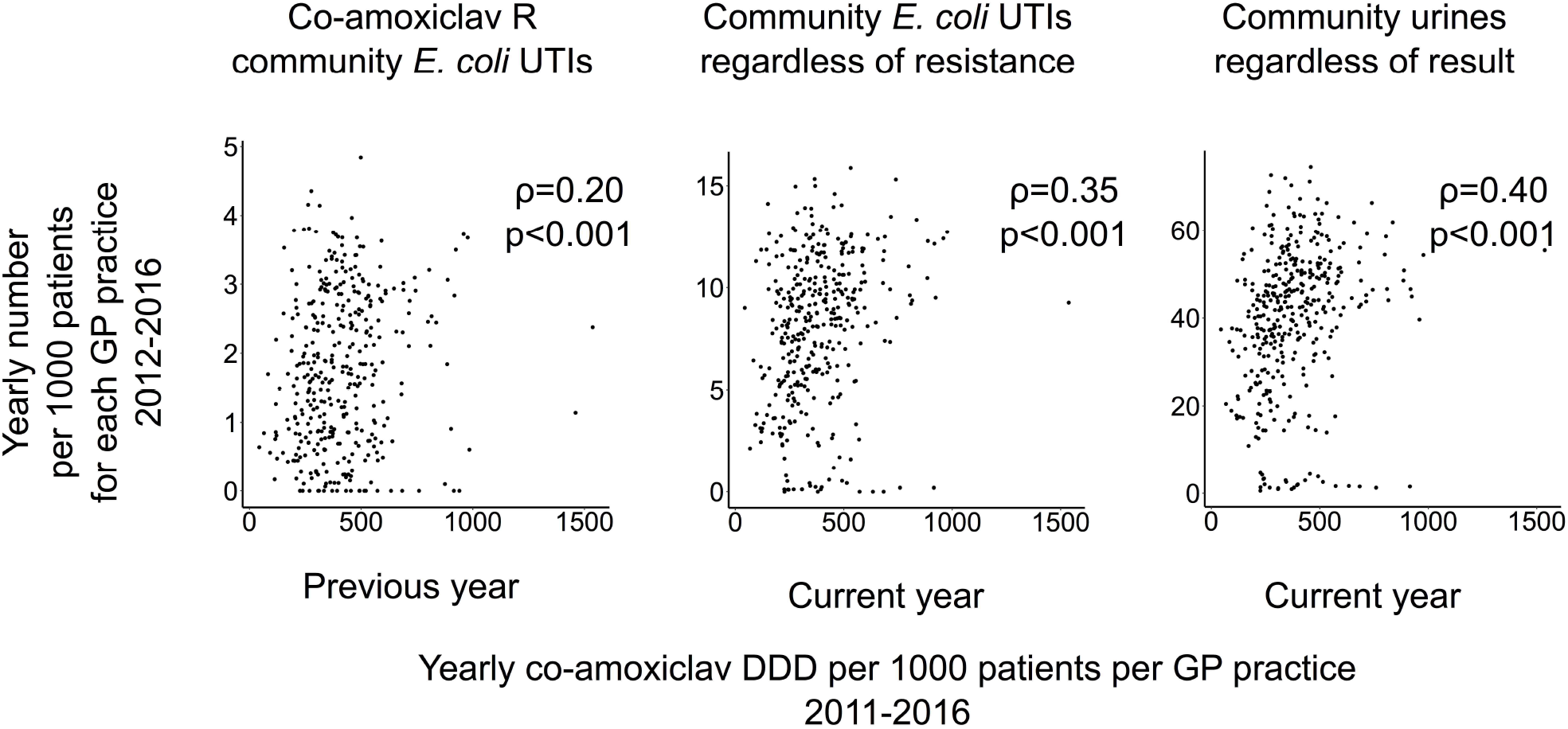
Number of community co-amoxiclav-resistant EC-UTIs (A), community EC-UTIs (B) and community urine samples (C) submitted regardless of result per 1000 patients per GP practice 2012-2016 compared with co-amoxiclav DDD per 1000 patients per general practice in the previous year for the first and the current year for the last two. Footnote: showing one record per year per GP practice. Spearman rho (and models) for each panel excludes 5 which submitted less than 151 samples over 2011-2016 (all others submitted over 308). Speraman rho for previous vs current for the 3 groups (ρ=0.20 vs ρ=0.04, ρ=0.33 vs ρ=0.35, ρ=0.37 vs ρ=0.40)

## Discussion

We have explored potential explanations for continuing increases in EC-BSI in Oxfordshire over 19 years using extensive, routinely-collected data, including diagnostic codes and laboratory/microbiology results. Incidence varied dramatically according to hospital-exposure, with increases notably being driven by community/quasi-community cases. This is important given a new National Health Service ambition aiming to reduce Gram-negative BSIs by targeting healthcare-associated cases; previous successful campaigns to reduce methicillin-resistant *Staphylococcus aureus* (MRSA) BSI and *Clostridium difficile* infections also focussed on nosocomial risk factors. Our data suggest that defining appropriate strategies targeting community/quasi-community associated EC-BSIs might have a greater impact. Crucially, co-amoxiclav-resistant EC-BSIs rose significantly faster than co-amoxiclav-susceptible EC-BSIs, regardless of hospital-exposure, with the greatest number of co-amoxiclav-resistant EC-BSIs in 2016 being community/quasi-community EC-BSIs. The association between primary care co-amoxiclav prescribing and co-amoxiclav-resistant EC-UTIs implicates co-amoxiclav prescribing as a key driver behind these rises. Co-amoxiclav is one of the most commonly prescribed antibiotics nationally in both the community and hospitals.^16,22^ Our findings indicate that reduced prescribing of co-amoxiclav could reduce the selection pressure for EC-BSI. Despite co-amoxiclav being used for empiric BSI treatment, there were no clinically important changes in mortality.

EC-BSI is generally considered ‘community-acquired’ although the true apportionment to community- vs healthcare-associated categories remains unclear, and there are differing definitions of healthcare-associated BSI.^6,23^ By linking to previous hospital admissions, one major study strength is that we could identify that incidence trends for non-nosocomial EC-BSIs varied significantly by proximity to hospital-exposure. Blood sample submission also increased significantly, potentially increasing ascertainment of ‘mild’ cases. However, blood cultures are key to the assessment of unwell patients whenever infection is suspected, and there were no clinically important changes in EC-BSI-associated severity at presentation or mortality, despite substantially increasing incidence, suggesting major ascertainment bias is unlikely.

The increasing trend in nosocomial EC-BSI was significantly smaller than for community/quasi-community EC-BSI in Oxfordshire, as observed nationally.^8^ Multiple infection control interventions were rolled out in UK hospitals from 2005-2010^24,25^ in response to MRSA/C. *difficile*, and horizontal components of these initiatives could have contributed to these lower nosocomial rates. Consistent with this, increases in hospital-onset BSI caused by Gram-negative bacilli reversed after a MRSA Prevention Initiative was introduced in the US, while community-acquired incidence did not change.^26^

Epidemiological differences between *E. coli*, MRSA and *C. difficile* also highlight the potential need for different interventions, particularly in primary care.^6^ In particular, recurrences explain relatively little of the ongoing increases in EC-BSIs, and both co-amoxiclav-resistant and co-amoxiclav-susceptible EC-BSI are rising. Overall, 42% of EC-BSI appeared to be more likely amenable to urinary-focussed intervention, similar to a England-wide study that found 51% of EC-BSIs had an underlying urogenital tract focus, with the largest independent risk factor for these being treatment for UTI in the prior four weeks.^8^ In our study, 13% of EC-BSIs were likely urine-associated and 28% presented as urosepsis; the first group may be most tractable for prevention but was smallest in both community and quasi-community EC-BSI, whereas urosepsis was the largest. One limitation is lack of data on visits to general practice; therefore, some patients may have had UTI symptoms and been treated empirically without a urine culture being sent, although successfully treated UTIs should not cause bacteraemia. Guidelines recommend urine samples be submitted from individuals with clinical treatment failure, frequent or recurrent UTI or with a possibly resistant infection;^16^ therefore bacteraemias due to UTI treatment failure should be ascertained within our data. It is hypothesised that much of the burden of EC-BSIs, and especially the rising incidence (in all hospital-exposure groups), arises from poor urinary catheter care. However, only 20% and 30% of the community and quasi-community groups, where incidence is increasing fasted, had a previous CSU, and there was no evidence that incidence was increasing faster in those with a previous CSU versus without. One key limitation is that we did not have records of the presence of a catheter, but only urine specimens recorded as being taken from a catheter, arguing that if a catheter was present and causing infection, a specimen would likely have been taken from it at some time.

Interestingly, there was strong evidence that quasi-nosocomial EC-BSIs with UTI or infectious diagnostic codes in the previous admission were rising faster than those without. This may reflect underlying predisposition to infection (e.g. chronic illnesses), or that prior antibiotic use adversely affects a patient’s microbiota potentially leading to colonisation/overgrowth by more pathogenic *E. coli*, thus predisposing to EC-BSI.

A limitation of surveillance studies is changes in antimicrobial susceptibility testing methodology (here in February 2013). Whilst the proportion of isolates classified as resistant can vary by testing protocol,^27,28^ crucially changes in co-amoxiclav resistance around this time occurred regardless of method (Supplementary Figure 14). Recent data suggest that broth dilution (BD-Phoenix) and the gold standard agar dilution have high agreement;^29^ thus, rising rates of co-amoxiclav-resistant (as defined by EUCAST breakpoints) EC-BSI/EC-UTI are likely correct.

For the first time, we have shown that GP practices with higher co-amoxiclav prescribing rates were more likely to have patients diagnosed with co-amoxiclav resistant EC-UTIs. Similar associations between trimethoprim use and trimethoprim-resistant urine-associated EC-BSI have been reported in adult women in England.^15^ Assessing usage-resistance associations is complicated, since changes in use of one antibiotic are generally accompanied by compensatory prescribing, and may be compounded by multi-drug resistance. Our results may therefore not be generalizable; for example, although the region we studied is sizeable (~1% of the UK), we did not observe a uniform decrease in cephalosporin-resistant and quinolone-resistant EC-BSIs as seen in BSI caused by Enterobacteriaceae.^14^ Such differences likely reflect a complex interplay of selection pressures.

A key limitation is that co-amoxiclav-resistant EC-BSI were too few over the period with contemporary prescribing data to investigate associations with antibiotic prescribing within the community. We were unable to assess associations between individual-patient antibiotic use (not available in the research database) and risk of resistant infections or between specific empiric regimens and outcome. However, there were no clinically important changes in mortality overall, by co-amoxiclav-susceptible/resistant phenotype, or by hospital-exposure across the study period. Co-amoxiclav remains our recommended first-line empiric treatment for most severe infections, so the substantial increase in incidence of co-amoxiclav-resistant bacteraemias suggests either that initial inappropriate treatment can be successfully rescued,^30^ or that the current definition of co-amoxiclav breakpoints may be suboptimal.^31^ Crucially, neither scenario supports a move towards broader empiric antibiotic treatment, consistent with prevailing antimicrobial stewardship messages.

In summary, on-going increases in EC-BSI were driven by community and quasi-community cases, and cannot be attributed only to increased recurrences or an aging population. Absence of changes in mortality and severity do not support ascertainment bias playing a major role, although this cannot be excluded. Whilst urinary foci are clearly important, at present the scope for intervening to prevent UTIs progressing to bacteraemia could be limited. Notably, higher co-amoxiclav use in primary care was associated with higher rates of both EC-UTI and co-amoxiclav-resistant EC-UTI, supporting drives to reduce broad-spectrum and inappropriate antibiotic use. However, despite substantial increases in co-amoxiclav-resistant EC-BSI, evidence that patient clinical outcomes are no worse does not support broadening empiric antibiotic prescribing from co-amoxiclav.^9^

## Acknowledgements

This work uses data provided by patients and collected by the NHS as part of their care and support. We thank all the people of Oxfordshire who contribute to the Infections in Oxfordshire Research Database. Research Database Team: R Alstead, C Bunch, DCW Crook, J Davies, J Finney, J Gearing (community), H Jones, L O’Connor, TEA Peto (PI), TP Quan, J Robinson (community), B Shine, AS Walker, D Waller, D Wyllie. Patient and Public Panel: G Blower, C Mancey, P McLoughlin, B Nichols.

Financial support: The research was funded by the National Institute for Health Research Health Protection Research Unit (NIHR HPRU) in Healthcare Associated Infections and Antimicrobial Resistance at the University of Oxford in partnership with Public Health England (PHE) [HPRU-2012-10041], and the NIHR Oxford Biomedical Research Centre, and a Medical Research Council UK Clinical Research Training Fellowship to NJF. The views expressed are those of the author(s) and not necessarily those of the NHS, the NIHR, the Department of Health or PHE. DWC and TEAP are NIHR senior investigators.

Contributions: KDV, NS, DHW, TEAP, and ASW designed the study. TPQ prepared extracts from the IORD database, KDV obtained data from the HSCIC. KDV and ASW analysed the data. KDV, TEAP and ASW prepared the figures. KDV, NS, and ASW prepared the first draft of the manuscript. All authors commented on the data and its interpretation, revised the content critically and approved the final version.

